# Identification and evolution of Cas9 tracrRNAs

**DOI:** 10.1101/2020.09.02.279885

**Authors:** Shane K. Dooley, Erica K. Baken, Walter Moss, Adina Howe, Joshua K. Young

## Abstract

Cas9 trans-activating CRISPR RNAs (tracrRNAs) form distinct structures essential for target recognition and cleavage and dictate exchangeability between orthologous proteins. As non-coding RNAs that are often apart from the CRISPR array, their identification can be arduous. In this paper, a new bioinformatic method for the detection of Cas9 tracrRNAs is presented. The approach utilizes a co-variance model (CM) based on both sequence homology and predicted secondary structure to locate tracrRNAs. This method predicts a tracrRNA for 98% of CRISPR-Cas9 systems identified by us. The identified tracrRNAs exhibit wide variation in sequence identity, however, CM analyses allow 94.7% to be categorized into just 10 related groups. Finally, association between Cas9 amino acid sequence-based phylogeny and tracrRNA secondary structure is evaluated, revealing strong evidence that secondary structure is evolutionarily conserved among Cas9 lineages. Altogether, our findings provide insight into Cas9 tracrRNA evolution and efforts to characterize the tracrRNA of new Cas9 systems.

## Introduction

CRISPR (clustered regularly interspaced palindromic repeats) RNA (crRNA) and CRISPR-associated (Cas) proteins cooperate to defend prokaryotic organisms against invading RNA and DNA.^1,2^ The Cas9 protein from type II CRISPR-Cas systems are guided to cleave double-strand (ds)DNA targets using two non-coding (nc)RNAs, a crRNA and a tracrRNA (trans-activating crRNA).^3,4^ The crRNA contains a sequence, termed the spacer, that directly base pairs with the dsDNA target site in the vicinity of a protospacer adjacent motif (PAM).^5–7^ The tracrRNA base pairs with the crRNA and is recognized and bound by Cas9 resulting in the formation of a dual guide RNA (gRNA) ribonucleoprotein (RNP) complex.^8,9^

In recent years, due to its RNA-based programmability, CRISPR-Cas9 has been widely adopted as a genome editing tool for a variety of different genomes including those from eukaryotic organisms.^8,10,11^ For these applications, the repair of a Cas9 induced double-strand break (DSB) has been harnessed to correct disease-causing mutations, introduce beneficial modifications (e.g. plant grain yield), and construct new biosynthetic pathways.^12–16^ To further simplify its use, the dual gRNA has been engineered into a single gRNA (sgRNA) by linking the crRNA and tracrRNA.^8^ Modifications to the Cas9 protein itself have also been made. By fusing new proteins domains to it and impairing its nuclease activity, it has been used as a robust RNA-guided DNA-binding platform. These applications include gene transcriptional activation and repression, epigenomic alteration, and precise DNA target deamination and modification.^17–24^

In prokaryotes, thousands of Cas9s have been identified computationally.^25–28^ In contrast, the gRNA solution for orthologous Cas9s may not be easily recognizable. This is mainly due to large variation in tracrRNA location, size and sequence identity.^27,29^ Consequently, the identification of tracrRNAs represents a bottleneck for the characterization of new Cas9 proteins and their development as genome editing tools. To address this limitation, several approaches have been developed. These include computational methods that locate tracrRNAs by using the CRISPR repeat sequence to search for the sequence in the tracrRNA that has homology to and base pairs with the crRNA (the anti-repeat). This is followed by a search for a Rho-independent-like termination signal in the vicinity of the anti-repeat. Other approaches reliant on the sequencing of the small ncRNAs transcribed from the CRISPR-Cas9 locus have also been used.^27,30^

Here, an algorithm was devised that combines previous methods with searches based on sequence homology and secondary structure co-variant models (CMs) to identify Cas9 tracrRNAs. Using this approach, a tracrRNA was located for greater than 98% of all CRISPR-Cas9 containing assemblies identifiable by us. Moreover, based on both sequence and structural homology, 94.7% of identified tracrRNAs could be unified into only 10 groups. Finally, Bayesian and non-parametric approaches quantifying phylogenetic signal revealed a strong evolutionary association between the Cas9 phylogeny and the predicted secondary structure of the tracrRNA, confirming previous observations that tracrRNA structures are a main determinant of Cas9-gRNA compatibility.^29,31^

## Materials and Methods

All custom code, scripts, parsers, python objects, and Jupyter Notebooks can be found on the primary author’s GitHub repository (https://github.com/skDooley/TRACR_RNA).

### Detection of CRISPR-Cas9 systems

Bacterial and archaeal assemblies were downloaded from PATRIC2, NCBI GenBank and RefSeq (last downloaded on May 05, 2020). CRISPR arrays were identified using MinCED v0.3.2 and PilerCR v1.06 with relaxed parameter settings (3 or more crRNAs, repeat lengths between 16 and 64 base pairs, and max spacer lengths of 64 base pairs).^32,33^

Next, a Hidden-Markov model (HMM) was generated from 83 previously described diverse Cas9 proteins using HMMER 3.2.1.^27,34^ The HMM was then used to search for Cas9-like proteins encoded in assemblies containing a CRISPR array. Protein sequences for each assembly were generated by translating open reading frames (ORFs) using Biopython to generate and filter ORFs for sequences between 673 and 2100 amino acids (Figure 1).^35^ Next, using BLAST, assemblies duplicated in our collection were removed.^36^

**Figure 1.**
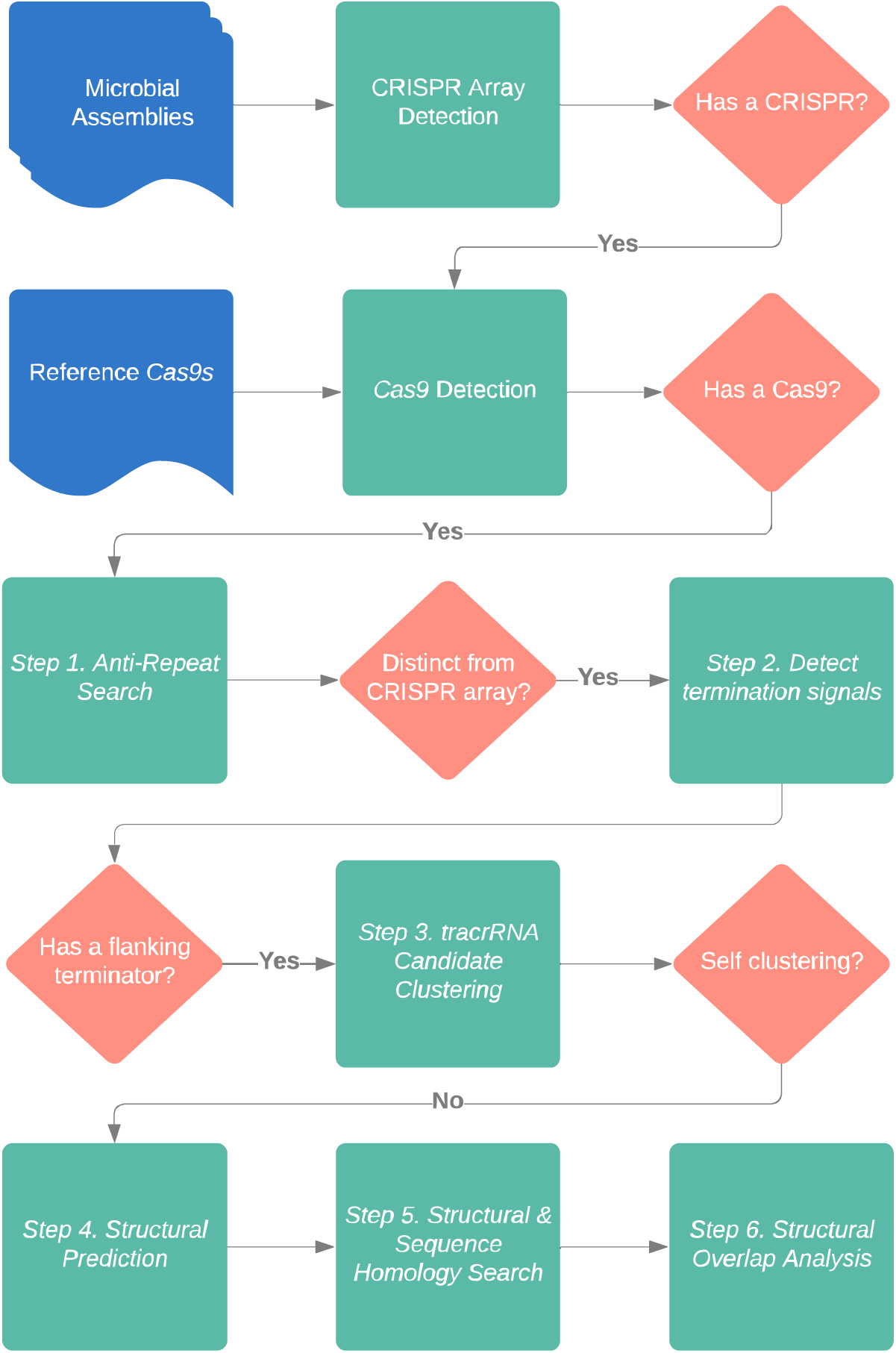
Cas9 tracrRNA detection pipeline. Flowchart of the informatic steps and key decisions points used to predict Cas9 tracrRNAs. Cas9-containing CRISPR systems are first identified in microbial DNA assemblies. Assemblies with a CRISPR-Cas9 loci are then searched in 6 steps to predict a tracrRNA. Inputs are shown in blue, informatic activities indicated in green and key decision points highlighted in orange.

The remaining assemblies and their Cas9 homologs’ were further examined for the presence of RuvC and HNH cleavage domains that define a Cas9 nuclease.^8,9^ This was initially accomplished through the visual inspection of protein alignments performed using MUSCLE between 83 diverse Cas9s described earlier for the key catalytic amino acids defining RuvCI, II and III subdomains and the HNH domain.^27,37^ Next, the identified regions were extracted and used to generate domain specific HMMs using HMMER 3.2.1. Each putative Cas9 protein from our collection was then scanned with the cleavage domain-specific HMMs. Proteins missing either domain or that had subdomains that were positional outliers were removed. Outlier determination was made by assessing the position of the RuvC I subdomain near the N-terminus and then comparing the relative distance of all other cleavage domains. Anything outside of three standard deviations (distribution of all the search results for the RuvC I subdomain) was removed except for the RuvC III subdomain, where proteins with more than four standard deviations from the mean distance were removed.

For phylogenetic signal analysis, the translated sequences were clustered at 90% sequence homology using CD-HIT v4.7.^38^ Representative sequences within each cluster were then selected and subsampled for calculating Cas9 and tracrRNA phylogenetic signal.

### Identification of Cas9 tracrRNAs

#### Step1: Search for anti-repeat signatures

The region of the tracrRNA capable of base pairing with the crRNA, the anti-repeat, was identified in Cas9-containing assemblies by searching for sequences with homology to the CRISPR repeat (using BLAST 2.7.3) that were in a region distinct from the CRISPR array (Figure 1: Step 1). While all assemblies had CRISPR arrays, neither of the two programs (PilerCR or MinCED) accurately detected all of the CRISPR repeats. To correct this and significantly reduce false positives, the coordinates of putative anti-repeats in the locus were referenced and used to identify locations that were at least one repeat-spacer unit length away from the CRISPR array.

#### Step 2: Detect Rho-independent termination signals

Rho-independent-like termination signals (RTS) were detected using ERPIN v5.5 (parameters -add 1 4 1 and -cutoff 100%) and an RTS database (Figure 1, Step 2).^39^ For this, the up- and down-stream regions adjacent to the anti-repeat were scanned for the presence of an RTS. Initially, each anti-repeat candidate with its respective RTS was considered a viable tracrRNA candidate. Additionally, if an anti-repeat had a termination signal on both sides, the pair were considered as potential tracrRNAs. Next, all tracrRNA candidates were conservatively filtered by removing sequences whose combined length was greater than 300 base-pairs. This cutoff-value was based on the longest characterized Cas9 tracrRNA length plus a generous buffer.^25^

#### Step3: Clustering tracrRNA candidates

Putative tracrRNAs were next clustered at 95% sequence identity with a 90% sequence coverage cutoff using cd-hit-est v.4.7.^38^ Sequence clusters that did not map back to at least five different assemblies were removed from further analysis unless they contained a putative tracrRNA from an assembly with only a single putative candidate (Figure 1, Step 3). The resulting sequences and their respective clusters then formed the basis for structural predictions.

#### Steps 4 and 5. tracrRNA structural predictions and searches for orthologous sequences

To generate a consensus secondary structure for each sequence-based tracrRNA cluster, sequences from each cluster were first aligned using MAFFT (--maxiterate 1000 --globalpair) and then fed into RNAalifold 2.4.5 (Figure 1, Step 4).^40^ The resulting consensus folds and sequences were then used as co-variance models within INFERNAL 1.1.2 to find RNA orthologs within the Cas9 associated DNA assemblies identified earlier (Figure 1, Step 5).^41^ All results of the CM search were then filtered to remove any hits whose corresponding nucleotide sequence had less than 55% pairing with either the consensus repeat or the reverse complement of the repeat.

#### Step 6. Analysis of CM overlap

Following tracrRNA identification, a final analysis was performed to examine the overlap between CMs. For this, CMs from steps 4 and 5 were used with INFERNAL 1.1.2 to identify similarities between each putative tracrRNA sequence cataloged in Steps 1-5. Results were next visualized by creating an undirected graph. In the graph, CMs were represented as vertices and a line was added between the two vertices if the CMs identified the same putative tracrRNA sequence. Connecting line widths were scaled by the percentage of shared sequences (percent similarity=(# of shared sequences)/(min(# found with model 1,# found with model 2))). Each network was then pruned for lines separating weakly connected vertices in order to isolate highly similar clusters. For phylogenetic analyses, all clusters not associated with the top 10 most common structures were removed to make statistical calculations computationally feasible.

### Calculating phylogenetic signal

To estimate the degree to which tracrRNA secondary structure associations are evolutionarily conserved among Cas9 lineages, phylogenetic signal was quantified using both Bayesian and non-parametric approaches. For the Bayesian approach, ancestral states of tracrRNA secondary structures along the Cas9 phylogeny were estimated using maximum likelihood under an All-Rates-Differ model in the R package diversitree.^42^ Achieving convergence with the full dataset was unattainable due to the computational complexity of estimating transition rates with more than 10 discrete tracrRNA states. Thus, the original dataset was pruned to include only the 10 most common tracrRNA secondary structures (as described above). Subsequently, this dataset was subsampled to represent 25% of the pruned data (512 lineages) while preserving the 62 verified tracrRNA sequences.^43^ With the ancestral state estimates, phylogenetic delta was calculated using a time-continuous discrete-trait Markov chain models (2 chains, 100,000 iterations each, thinned every 10 iterations, 100 iterations deleted as burn-in, see Borges et al. 2019 for more details). Values of phylogenetic delta above 1 indicate a close correspondence of the trait with the phylogeny, with increasing values representing increasing correspondence (i.e., strong phylogenetic signal), whereas values near 0 indicate weak phylogenetic signal. The results presented below were generated from a subsampling procedure that successfully converged. The Cas9 lineages involved in this calculation can be found in the supplemental materials (Supplementary Table S1). To ensure the results were not biased by subsampling, ten iterations were performed, and the resulting delta values for each round of subsampling can be found in the supplemental materials (Supplementary Table S2).

The second approach for quantifying phylogenetic signal was a modified two-block partial least squares test.^44^ This procedure utilized the pruned dataset described above prior to the resampling procedures (2050 lineages included) and quantified the correlation coefficient of the Cas9 phylogeny (converted to a phylogenetic covariance matrix) with the trait matrix. Multivariate effect size and significance were calculated using residual randomization via permutation procedures (1000 iterations, R package geomorph).^45,46^

## Results

### Identification of type II CRISPR-Cas9 systems

41,999 putative type II CRISPR-Cas9 systems from over 1 million microbial nucleotide sequences were identified (Supplementary Table S3). Cas9 length ranged from 700 to 1,800 amino acids and exhibited a bimodal length distribution centered around 1,100 and 1,400 amino acids (Supplementary Figure S1). To calculate the phylogenetic relationship between tracrRNA and Cas9, 2,724 diverse and representative systems were also selected (Supplementary Table S3) and subsampled (Methods section and Supplementary Table S1). Additionally, as a control for our methods, 79 type II CRISPR-Cas9 systems with an experimentally established tracrRNA were also included in our analysis (Supplementary Table S4).^43^ Of these, a Cas9 encoding ORF was detected for 73 using our methods.

### Co-variant detection of Cas9 tracrRNAs

The detection of Cas9 tracrRNAs was automated using a multi-step approach that combines both homology and structural searches (Figure 1). First, taking advantage of previous methods, the identification of the tracrRNA anti-repeat and Rho-independent termination-like signal were automated similar to those described in Chyou *et al*., 2019 (Figure 1, Steps 1 and 2).^27,47–49^ Next, based on functional associations between gRNA secondary structure and orthogonality, we reasoned that conserved tracrRNA structural features could be used to complement homology-dependent methods in the identification of a tracrRNA.^31^ To accomplish this, sequences of tracrRNAs predicted in Steps 1 and 2 (Figure 1) were first aligned and clustered based on sequence similarity (Figure 1, Step 3). Next, sequences within each cluster were used to predict a consensus secondary structure (Figure 1, Step 4). CMs were next generated from each cluster based on sequence and structural homology and used to search for related tracrRNAs (Figure 1, Step 5). Finally, to examine the relationship between tracrRNAs in our collection, a last clustering step based on CM similarity was applied (Figure 1, Step 6). For some systems, multiple solutions were observed within the CRISPR-Cas9 locus after Step 6 (Figure 1) (Supplementary Figure S2A-E). In these cases, additional filtering was applied to permit the selection of a single tracrRNA. First, the tracrRNA that was closest to the *cas9* gene was selected. If two or more tracrRNAs in the region closest to the *cas9* gene had overlapping locations, the tracrRNA with the most stable secondary structure (based of minimum free energy calculations of the predicted folds) was chosen.

Next, our pipeline was used to predict tracrRNA solutions for the 41,999 CRISPR-Cas9 systems identified earlier. To establish false positive and negative rates of our approach, its ability to accurately predicted the tracrRNA from a curated set of 74 experimentally proven Cas9 tracrRNAs was also evaluated.^43^ For this, the loci containing the curated tracrRNAs were identified and flagged in our collection. Altogether, our algorithm predicted a tracrRNA for 98% (41741 out of 41999) of the Cas9 systems searched (Supplementary Table S3). For the curated set of tracrRNAs proven to support Cas9 functionality, our approach correctly identified 90.4% (66 out of 73) resulting in false positive and negative rates of 6.8% (5 out of 73) and 2.7% (2 out of 73), respectively. Of the five systems where a different tracrRNA was identified, four (Cco, Kki, Lsp2 and Nsa) were predicted to have a tracrRNA that was transcribed in the opposite direction from the anti-repeat than described earlier. For the fifth system, the tracrRNA was predicted to be in a different location (Ghy3) (Supplementary Figure S2A-E).^43^

### Sequence and structural homology of Cas9 gRNAs

Based on sequence and structural overlap, 94.7% (39527 out of 41741) of identified tracrRNAs could be categorized into 10 clusters (Figure 2). 1,388 of the 2,214 remaining tracrRNAs were classified into 31 additional CM-based similarity groups and 826 of the remainders represented as singletons in the dataset (Supplementary Table S3). The majority of previously characterized tracrRNAs could be found in the 10 most abundant clusters (Figure 2, Clusters 1-7 and 10).

**Figure 2.**
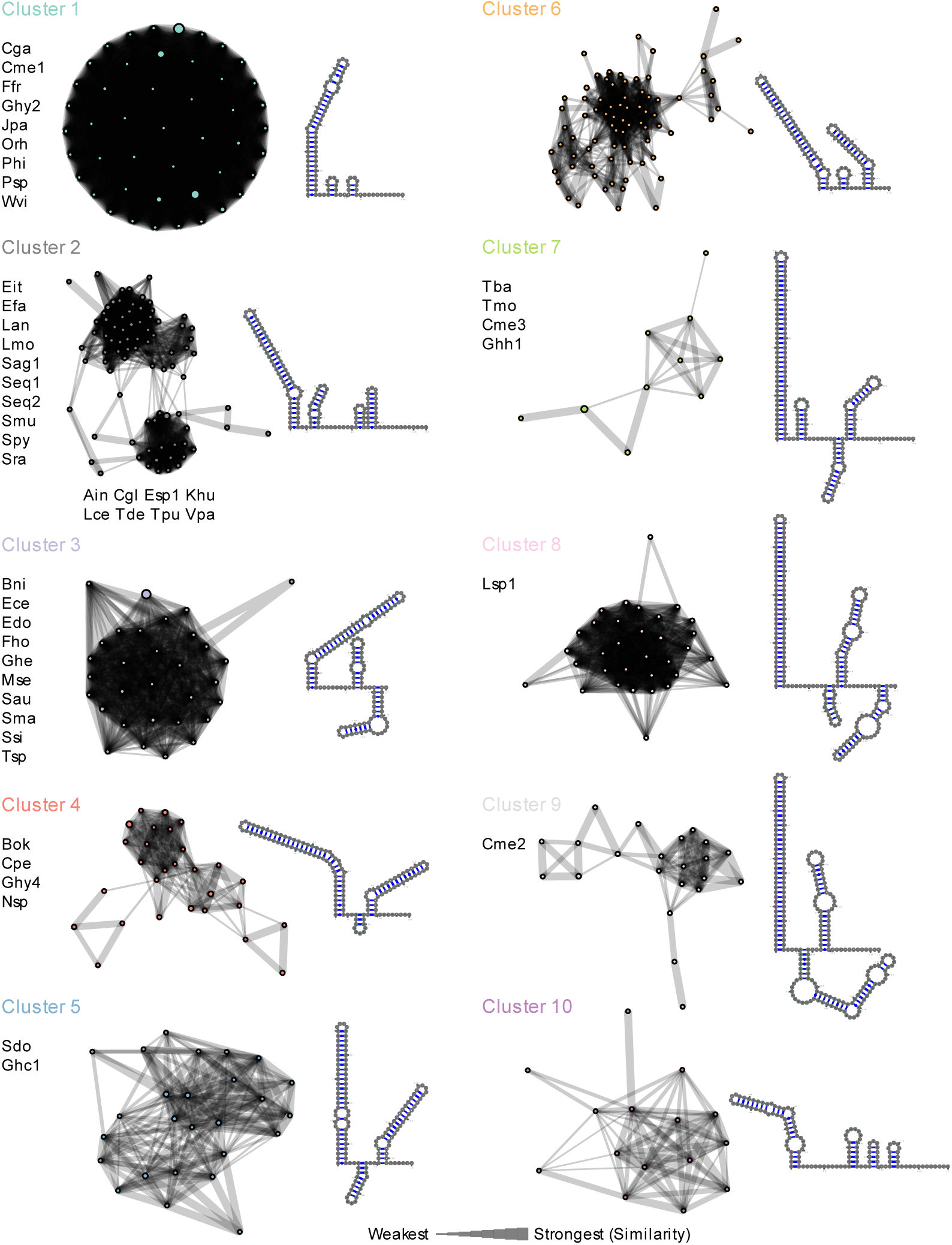
Top 10 co-variant models and clustering of Cas9 tracrRNAs. Top 10 Cas9 tracrRNA clusters based on similarity between sequence and predicted secondary structure co-variant models (CMs). Circles represent a CM and are colored according to the designated cluster. The width of the connecting lines indicates the percentage of similarity or relatedness among CMs. Previously characterized tracrRNAs associated with each cluster are indicated. Single guide RNA (sgRNA) solutions for the most abundant tracrRNA sequence are also shown immediately adjacent to each cluster. The lower stem, bulge, upper stem and nexus regions are colored purple, orange, teal, blue and green, respectively.

To visualize tracrRNA structural features in the context of the dual gRNA used by Cas9, a sgRNA was generated by linking the 3’ end of the full-length CRISPR repeat with a self-folding tetraloop (5’-GAAA-3’) to the 5’ end of the anti-repeat in the tracrRNA as described previously.^50^ This was done once for each of the top 10 clusters using the most abundant tracrRNA sequence and respective CRISPR repeat. As observed previously, most sgRNA structures comprised varying degrees of complementation between the repeat and anti-repeat followed by two or more hairpin-like structures in the tracrRNA (Figure 2).^29,31,43^ Likewise, a repeat:anti-repeat mismatch resulting in a bulge was detected in some but not all instances (Figure 2). In most cases, the nexus fold, a functionally important and conserved hairpin structure hypothesized to orient the spacer away from the rest of the dual gRNA, was detected almost immediately (within 2 or 3 nts) after the repeat:anti-repeat duplex (Figure 2, Clusters 1, 2, 4-6, 7 and 9)^25,31^. For clusters 3, 8 and 10, it was located approximately 9 nts after the repeat:anti-repeat (Figure 2). The nexus-like fold itself varied in length from 10-80 nts with an average length of 24 nts and ranged from simple 3 nt stem loop structures (Figure 2, Clusters 1, 4 and 6) to more complex structures with additional bulges and stems (Figure 2, Clusters 3 and 9).

### Cas9 and tracrRNA evolutionary association

Cas9 phylogeny and predicted gRNA secondary structures have been linked to exchangeability between orthologous Cas9s.^29,31^ This suggests tight evolutionary association between Cas9 and its gRNA structural features. To further test this observation, we examined the phylogenetic association between tracrRNA secondary structure and Cas9 protein. For this, two statistical methods, Bayesian estimation of the phylogenetic delta statistic and a non-parametric modified two-block partial least squares model, were used to evaluate the phylogenetic relationship between the 10 primary tracrRNA secondary structures (encompassing 83.3% of all representative tracrRNAs identified) and our representative collection of Cas9 proteins. First, a diverse and representative collection of Cas9s were subsampled and a phylogenetic tree was constructed. Next, tracrRNA structures were mapped to it (Figure 3). Ancestral states were then estimated, from which the delta statistic was calculated. In these scenarios, a significant phylogenetic signal was detected using both approaches (delta = 244.107 and r-PLS = .912, effect size = 28.952, p =1e-04) as can be visualized by the strong clustering of tracrRNA secondary structures across Cas9 phylogeny (Figure 3). Rare exceptions to this were observed as the occurrence of the same tracrRNA structure in distantly related orthologs (Figure 3).

**Figure 3.**
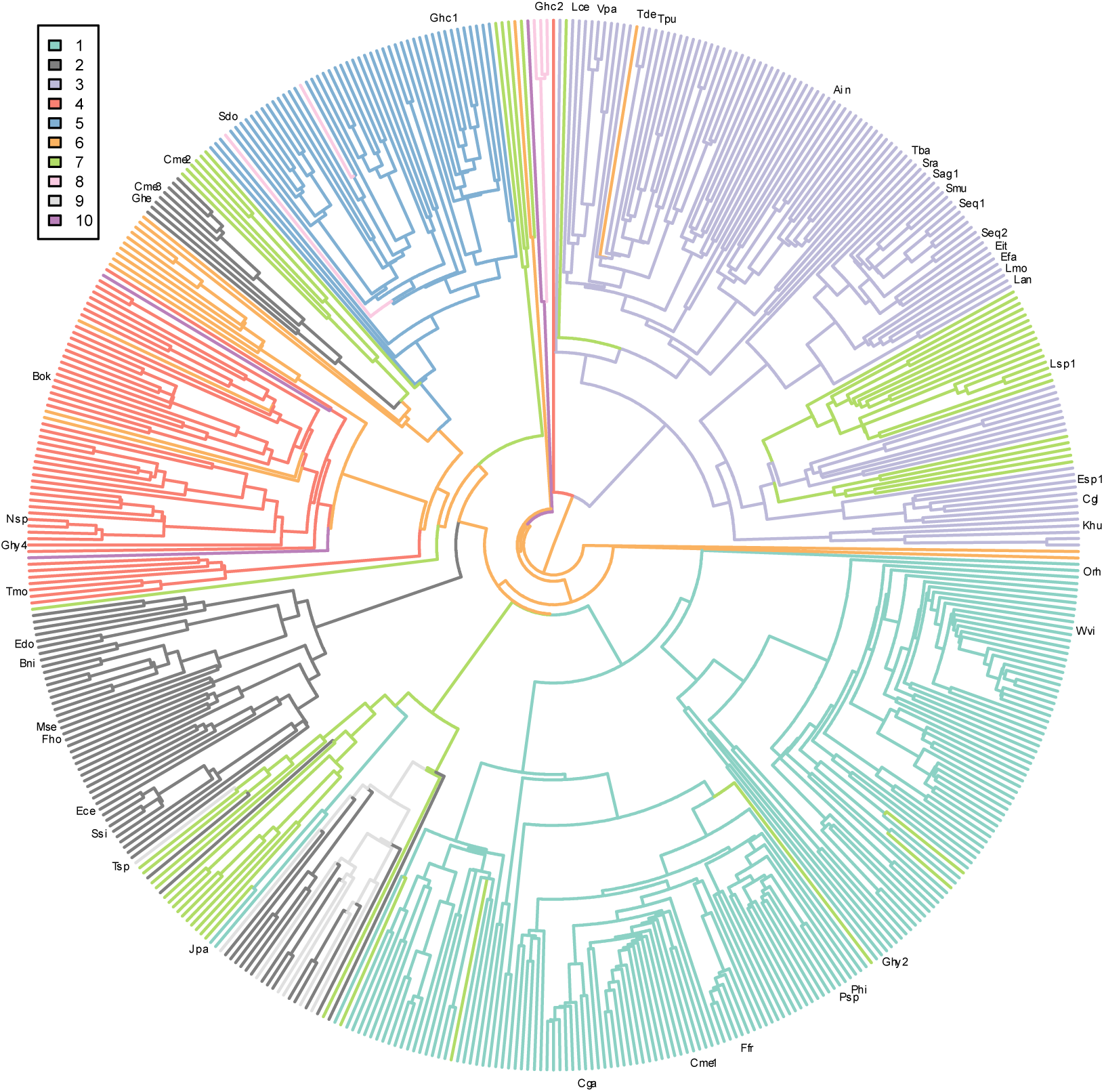
Cas9 phylogeny and tracrRNA secondary structure. Predicted tracrRNA secondary structures associated with Cas9 phylogeny. Each color represents tracrRNA secondary structure associated with Clusters 1-10 (Figure 2). Cas9 proteins characterized previously are indicated (Supplementary Table S4).

## Discussion

We provide a framework for the global identification of CRISPR-Cas9 tracrRNAs. Our method builds upon previous approaches ^27,47–49^ that have sought to identify the key components that define a tracrRNA, the anti-repeat, and 3’ hairpin-like secondary structures, and adds to them by utilizing CMs to identify sequence and structural homologs (Figure 1, Steps 1-5). In total, we predicted a tracrRNA solution from 98% of the identified Cas9 systems. In comparison with a diverse collection of experimentally determined tracrRNAs, we also showed that our approach in most cases (66 out of 73 (90.4%)) could accurately identify a Cas9 tracrRNA.^43^ In the five instances where a different tracrRNA was detected (Cco, Ghy3, Kki, Lsp2 and Nsa), it is possible that a second tracrRNA may have evolved in the CRISPR-Cas9 locus as described previously.^25^ This is supported in part by the identification of alternative tracrRNAs in these loci that exhibit CM homology to tracrRNAs known to support Cas9 functionality (Figure 2 and Supplementary Figures S2A-E). Additionally, the location of the alternate tracrRNA within the CRISPR-Cas9 locus is consistent with other characterized systems. These locations include regions near the end of the CRISPR array or directly adjacent to the *cas9* gene (Supplementary Figures S2A-E).

In examining the sequence and structural overlap of the identified tracrRNAs using CMs (Figure 1, Step 6), we found that they could be classified mainly into 10 groups (Figure 2). Interestingly, when observing the distribution for previously determined tracrRNAs, it seems that our structural classifications also correlated with Cas9-gRNA compatibility (Figure 2).^29,43,51^ Although, in some cases, as observed in cluster 2, Cas9 and gRNAs (Spy and Tde) previously shown to be incompatible were grouped together (Figure 2).^51^ This finding indicates that features in the repeat:anti-repeat duplex may also confer non-compatibility or that some clusters represent a continuum of related tracrRNA structures that diverge into non-compatible ones. Indeed, the latter point seems more likely for Spy and Tde provided the related, yet almost distinct groupings observed within cluster 2 (Figure 2). Altogether, our findings suggest the number of non-cross reactive gRNA groupings may be extended from 7 to 10 or more.^31,43,51^ Furthermore, this finding expands the number of potential Cas9s available for orthogonal genome editing approaches or applications that require RNA-guided multiplexing.^51,52^

Both approaches for calculating phylogenetic signal showed that the tracrRNA structure is an evolutionarily conserved trait among Cas9 lineages. This matches previous observations that Cas9-gRNA exchangeability is associated with gRNA secondary structure and Cas9 phylogeny.^29,31^ Interestingly, exceptions to this were noted in our analysis. In those instances, distantly related Cas9 orthologs were associated with the same tracrRNA structural classification. This observation may provide evidence of more recent evolutionary events resulting from the resetting of the tracrRNA or recombination between different CRISPR-Cas9 systems.^25^

## Conclusion

Using CMs based on both sequence homology and predicted structure, an informatic approach, enabling the identification of Cas9 tracrRNAs, was developed. This method permitted the global identification of more than 41K tracrRNAs and the development of sgRNA solutions for nearly all CRISPR-Cas9 systems detected by us. Structural predictions revealed strong homology among tracrRNA secondary structure that tightly correlated with the Cas9 phylogeny allowing the majority of Cas9 tracrRNAs to be classified into just 10 groups. Altogether, the results presented here will aid in the characterization and development of new Cas9s as genome editing tools and maybe extended to other CRISPR systems that utilize a tracrRNA.^53–56^

## Acknowledgements

We thank Dr. Dean Adams (Iowa State University) for his advice and expertise on how to calculate phylogenetic signal and for his quick replies to bugs in the phylogenetic analysis. Additionally, thank you to Kevin Hayes for supporting the early stages of this analysis and Corteva for funding the initial stages of this research. We thank the ISU legal team and Lynne Mumm for her work in facilitating the collaboration agreements between Corteva and Iowa State. Finally, thank you to Dr. James Reece and the Office of the Vice President for Research at Iowa State University for funding me (SKD).

## Authorship Confirmation Statement

SKD, EKB, WM, AH and JKY designed research; SKD performed research and SKD, EKB and JKY analyzed data. SKD, EKB, AH, and JKY wrote the paper. All authors read and approved the final manuscript and it has not been published, in press, or submitted elsewhere.

