## Supplementary Figure S1 for "Identification and evolution of Cas9 tracrRNAs"

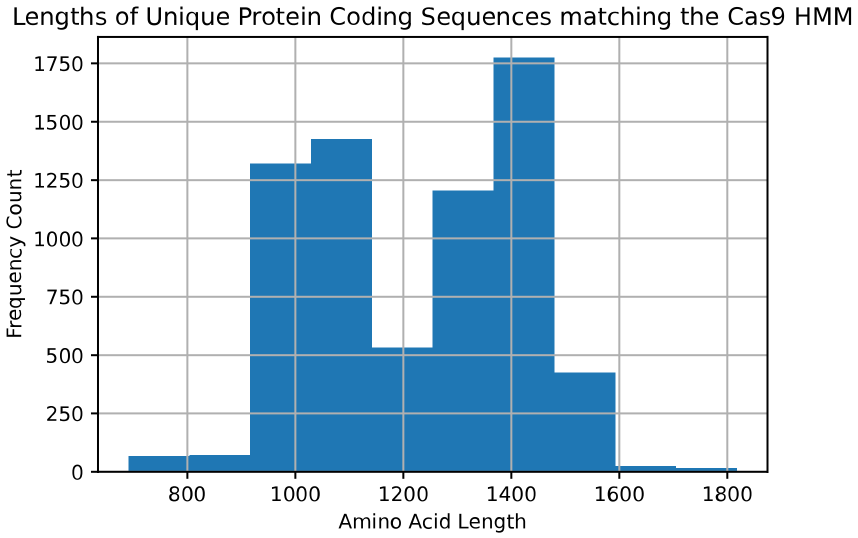


**Supplementary Figure S1**. Distribution of the coding-sequence lengths of the identified Cas9 proteins.
