## Supplementary Figure S2 for "Identification and evolution of Cas9 tracrRNAs"

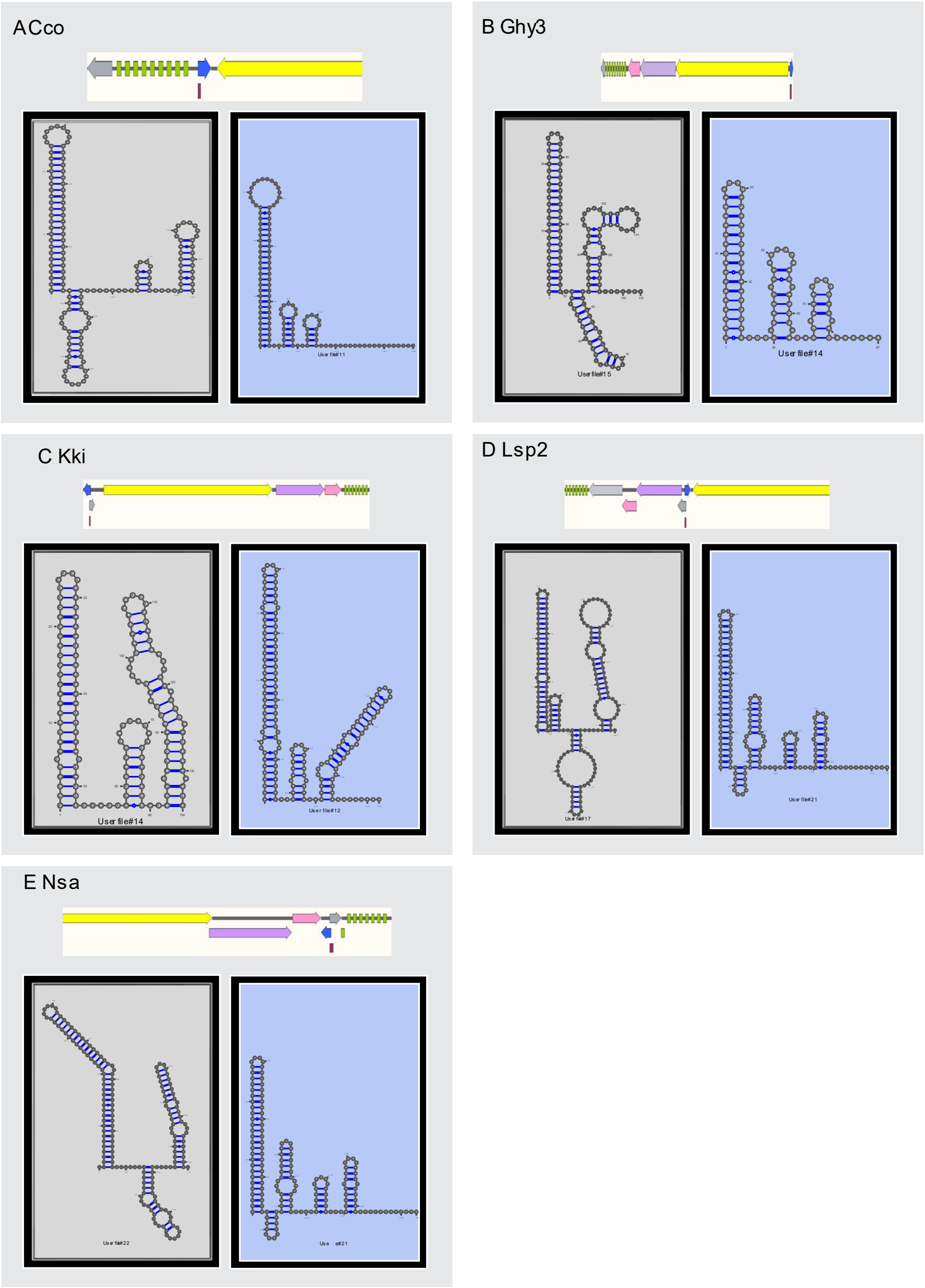


**Supplementary Figure S2**. CRISPR-Cas genomic locus annotations for *Campylobacter coli* (Cco) (**A**), Geyser-Hotspring_Yellowstone (Ghy3) (**B**), *Kingella kingae* (Kki) (**C**), Lsp2 (*Lactobacillus sp.*) (**D**), and *Nitratifractor salsuginis* (Nsa) (**E**). The predicted and experimentally determined tracrRNAs are in gray and blue, respectively. The *cas9* gene is in yellow, CRISPR repeats in green, *cas1* and *2* genes in purple and pink, respectively, and *csn2* gene (only present for Lsp2) in light gray. Arrows indicate the orientation of each gene or tracrRNA. Below each locus are the predicted sgRNA structure (gray) and the experimentally verified sgRNA (blue). In these five cases, the predicted sgRNA is formed from a tracrRNA with structural homology to experimentally verified tracrRNAs.
